# Serum level of High-density lipoprotein particles are independently associated with long-term prognosis in patients with coronary artery disease: The GENES study

**DOI:** 10.1101/676122

**Authors:** Thibaut Duparc, Jean-Bernard Ruidavets, Annelise Genoux, Cécile Ingueneau, Souad Najib, Jean Ferrieres, Bertrand Perret, Laurent O. Martinez

## Abstract

**BACKGROUND:** HDL-Cholesterol (HDL-C) is not an accurate surrogate marker to measure the cardioprotective functions of HDL in coronary artery diseases (CAD) patients. Hence, measurement of other HDL-related parameters may have prognostic superiority over HDL-C.

**OBJECTIVE:** This work aimed to examine the predictive value of HDL particles profile for long-term mortality in CAD patients. Its informative value was compared to that of HDL-C and apoA-I.

**METHODS:** HDL particles profile were measured by nuclear magnetic resonance (NMR) spectroscopy in 214 male participants with stable CAD (45-74 years). Median follow up was 12.5 years with a 36.4% mortality rate. Cardiovascular mortality accounted for 64.5 %.

**RESULTS:** Mean concentrations of total HDL particles (HDL-P), small-sized HDL (SHDL-P) and apoA-I were lower in deceased than in surviving patients whereas no difference was observed according to HDL-C and large HDL particles. All NMR-HDL measures were correlated between themselves and with other HDL markers (HDL-C, apoA-I and LpA-I). In a multivariate model adjusted for 14 cardiovascular risk factors and bioclinical variables, HDL-P and SHDL-P displayed the strongest inverse association with all-cause and cardiovascular mortality. Weaker associations were recorded for apoA-I.

**CONCLUSION:** HDL particle profile measured by NMR spectroscopy should be considered to better stratify risk in population at high risk or in the setting of pharmacotherapy.

## Introduction

HDL-Cholesterol (HDL-C) has been repeatedly inversely related to cardiovascular risk in all epidemiological studies. However, pharmacological trials aimed at increasing HDL-C have failed to demonstrate a beneficial effect on clinical outcomes^1^. Similarly, genetic variants associated to increased HDL-C have not been found associated to a decreased cardiovascular risk^2^. This has led to the concept that a single measurement of HDL-C does not necessarily reflect the functional properties of HDL particles and their effects against atherosclerosis. Indeed, HDL particles are heterogeneous in size and biochemical composition, and HDL subpopulations might have different functional properties^3^. NMR-spectroscopy has been recently proposed as a tool to quantify HDL particles and HDL subpopulations^3^. This technology enables to measure the total concentration of HDL particles and their size distribution. Numerous recent studies have shown that the atheroprotective properties of HDL are supported by small and medium-sized HDL particles^4^, which were inversely related to cardiovascular risk in various clinical settings^5,6^.

In the present study, we have evaluated HDL particles concentration and distribution in a cohort of patients with established, angiographically documented, coronary artery disease^7,8^ taking into account an extended panel of potential confounders related to cardiovascular risk and heart condition. The patients’ vital status was yearly assessed and mortality was recorded, distinguishing all-cause mortality, cardiovascular mortality and other causes of death, during a 12.5-year median follow-up. In addition, one objective of the study was to compare HDL particles measurements to routinely available HDL markers, HDL-C and apoA-I, as predictors of mortality in CAD patients. Indeed, apoA-I is the major HDL protein, its immunoassay is today referred to international standards and now available on automated analyzers. Moreover apoA-I is much less influenced than HDL-C by other components of the lipoprotein profile, like VLDL or LDL, which may have an impact on HDL lipid composition through action of lipid transfer proteins.

In this study, multivariate analyses demonstrate that total HDL particles (HDL-P), small-sized HDL (SHDL-P) and apoA-I are predictors of all-cause and cardiovascular mortality in coronary patients.

## Materials and Methods

### Study participants

The “Génétique et Environnement en Europe du Sud” (GENES) study is a case-control study designed to assess the role of genetic, biological and environmental determinants in the occurrence of CAD^9^. All participants signed an informed consent form. The study protocol was approved by the local ethics committee (CCPPRB, Toulouse / Sud-Ouest, file #1-99-48, Feb 2000). A biological sample collection has been constituted (declared as DC-2008-463 #1 to the Ministry of Research and to the Regional Health Authority). As previously described, cases were stable male CAD patients living in the Toulouse area (South-west France), aged 45-74 and prospectively recruited from 2001 to 2004 after admission to the Cardiology department, Toulouse University Hospital, for cardiovascular examination and referred for evaluation and management of their CAD^10^. Stable CAD was defined by a previous history of acute coronary syndrome, a previous history of coronary artery revascularization, a documented myocardial ischemia, a stable angina or the presence at coronary angiography of a coronary stenosis of 50% or more. Patients who had presented an acute coronary episode during the past eight days were not included in the study, because they were considered unstable. In the present analysis, we only took into account the first 214 consecutive patients in whom NMR-HDL profile was measured and complete data were available for all the subjects.

### Assessment of the vital status

Vital status on December 31, 2014, was obtained for each participant through the national database (“RNIPP”), which records, every year, all deaths occurring in the French population (http://cesp.vjf.inserm.fr/svcd). Vital status was assessed yearly, with a median follow up of 12.5 years. All dates and causes of death were obtained for participants who died during the follow-up. Main and associated causes of deaths were provided by the French National Institute of Health Research (CépiDc-INSERM), which systematically collects and codes (using the International Classification of Diseases coding system) data recorded on death certificates. Death from a cardiovascular cause during follow-up was assessed by a committee of four medical doctors, every time cardiovascular disease was reported as the main cause of death, or when it was mentioned as an associated cause, if the main cause was a plausible complication of CV disease. Authorizations to use these data were obtained in accordance with French law (Commission nationale de l’informatique et des libertés (CNIL): authorization 355152v1, September 3, 2008).

### Biological measurement

Blood was collected after an overnight fast. Serum sample aliquots were subsequently stored at - 80°C until biological analyses. The following biomarkers were assayed with enzymatic reagents on automated analyzers (Hitachi 912 and Cobas 8000, Roche Diagnostics, Meylan, France): serum total cholesterol, HDL-C, triglycerides, fasting glucose, creatinin. eGFR was calculated using the abbreviated Modification of Diet in Renal Disease (MDRD) Study equation^11^. ApoA-I, high-sensitive C-Reactive protein (hs-CRP), N-terminal pro-brain natriuretic peptide (NT-proBNP) and high-sensitive cardiac troponin T (hs-TnT) were determined on the same analyzer by immunoturbidimetry assays (Roche Diagnostics). Lipoprotein A-I (containing apoA-I but not apoA-II) were measured by immunoelectrodiffusion^8^.

### Data collection

Age, environmental characteristics and information on cardiovascular risk factors were collected through standardized face-to-face interviews, performed by a single physician. Past medical history was collected and checked in the patients’ medical files. Presence of dyslipidemia, diabetes mellitus or hypertension was assessed from the subjects’ current treatments. Dyslipidemia was defined as treatment with drugs or fasting serum total cholesterol ≥ 2.40 g/L. Hypertension was defined as treatment with drugs or systolic blood pressure ≥160 mmHg or diastolic blood pressure ≥ 95 mmHg. Diabetes was defined as treatment with drugs or fasting blood glucose ≥ 7.8 mmol/L. Smoking status was classified as current smokers, past smokers having quitted for more than 3 years and patients having never smoked. Among current smokers, cigarette consumption was estimated with the pack-year quantification and recorded as the average number of cigarettes per day. Alcohol consumption was assessed using a typical week pattern. The total amount of pure alcohol consumption was calculated as the sum of different types of drinks and was expressed as grams per day. Physical activity was investigated through a standardized questionnaire^12^ and categorized into three levels as: no physical activity, moderate physical activity during 20 minutes no more than once a week, and high physical activity during 20 minutes, at least twice a week. Blood pressure and resting heart rate were measured with an automatic sphygmomanometer (OMRON 705 CP). Measurements were performed after a minimum of 5 minutes rest; average values from two different measurements were recorded for further analysis.

### Assessment of CAD severity and extension and estimation of cardiac function

Coronary artery stenoses of ≥ 50% luminal narrowing were considered significant. Diffusion of coronary artery disease lesions was assessed by calculating the Gensini Score, based on data from coronary angiography^13–15^. Left Ventricular Ejection Fraction (LVEF) was assessed by contrast ventriculography using an isotopic method, and/or by echocardiography.

### HDL measurement by Nuclear Magnetic Resonance (NMR) spectroscopy

HDL particle concentration and size were measured by NMR spectroscopy using the AXINON^®^ lipoFIT^®^-S100 test system (Numares AG, Regensburg, Germany) as previously described^16–18^. Serum (630 μL) gently mixed with 70 μL of an additives solution containing reference substances, NaN3 and D2O, and 600 μL of the mixture were transferred into 5 mm NMR tubes with barcode-labeled caps. Briefly,^1^H NMR spectra were recorded at a temperature of 310 K on a shielded 600 MHz Avance III HD NMR spectrometer (Bruker Biospin) with a 5 mm triple resonance TXI probe head including deuterium lock channel, a z-gradient coil and automatic frequency tuning and matching. Prior to each analytical run, calibration was performed using a calibration sample comprising an aqueous solution of various calibration substances with different molecular masses, 0.01% (w/v) NaN3, 10 % (v/v) D_2_O as a locking substance and 1 % glycerol to adjust viscosity. Two identical control samples were measured directly after calibration and at the end of each run. Each spectrum was referenced, normalized and subjected to a set of quality checks including checks of baseline properties, noise level, shift, width, and symmetry properties of quality control signals. Lipoprotein analysis was conducted via deconvolution of the broad methyl group signal at about 0.9-0.8 ppm. In this process, lipoprotein subclasses are reflected by a fixed number of pre-defined bell-shaped (e.g. Gaussian or Lorentzian) base functions, each of which has a constant position and defined width. The concentrations of lipoprotein particle subclasses as well as the average particle size were calculated based on the integrals attributable to specific base functions. Fit quality was checked by calculating the residual deviation between fit and spectrum intensity. In this study, the concentrations of large-sized HDL particles (LHDL-P), small-sized HDL-particle (SHDL-P) and total HDL particles (HDL-P, reported in nmol / L) as well as the average HDL particle size (HDL size, reported in nm) are used. The two measured HDL subclasses had the following estimated diameter ranges: LHDL-P, 8.8 − 13 nm; SHDL-P, 7.3 − 8.7 nm.

### Statistical analyses

Continuous variables are displayed as means and standard deviations (SD). Categorical variables are presented as proportions. We first described and compared characteristics of participants according to vital status. Categorical variables were compared between groups using the χ^2^-test (or Fisher’s exact test when necessary). Student’s *t*-test was used to compare the distribution of continuous data. A Wilcoxon Mann-Whitney’s test (or logarithmic transformation of the variable when necessary) was performed when distribution departed from normality, or when homoscedasticity was rejected. Spearman rank correlations were used to test the associations of NMR-HDL parameters and HDL-C with cardiovascular risk factors, severity, extension and estimation of cardiac function of the disease.

Cumulative survival of patients were determined by the Kaplan-Meier method and compared, using the Log-rank test for the individual endpoints of all-cause mortality. The relation between baseline variables and mortality was assessed using Cox proportional hazards regression analysis. We tested the proportionality assumption using cumulative sums of martingale-based residuals. We performed regression analyses with polynomial models (quadratic and cubic) to examine for possible non-linear relations between continuous variables and mortality. Cox regression analyses were performed first without any adjustment for co-variables and, second, with adjustment on classical cardiovascular risk factors (age, smoking, treatments for dyslipidemia, hypertension and diabetes). Further adjustments were successively performed on extended cardiovascular risk factors (alcohol consumption, physical activity, BMI, eGFR, hs-CRP and duration of CAD) and clinical parameters related to the severity, extension of the disease and cardiac function (heart rate, LVEF and Gensini score). All statistical analyses were carried out using the SAS statistical software package 9.4 (SAS Institute, Cary, NC). Analyses were two-tailed and p < 0.05 was considered to be significant.

## Results

### Characteristics of CAD patients according to vital status

The present cohort was constituted of 214 CAD male patients. After inclusion, the vital patients’ status was yearly assessed. The median follow-up period was 12.5 years (mean: 10.7 years). During follow-up, 78 deaths had been recorded giving a death rate of 36.4 % and a mean annual rate of 3.4%. Cardiovascular mortality accounted for the majority of deaths recorded (64.1 %, n = 50) and cancers accounted for 16.7 % (n= 13).

Comparison of patients’ data when they were included in the cohort is given in Table 1. Results are presented distinguishing two groups: the ‘‘alive group’’ (patients living at the end of the follow-up) and the ‘‘deceased group’’ (patients who died during follow-up).

**Table 1.**
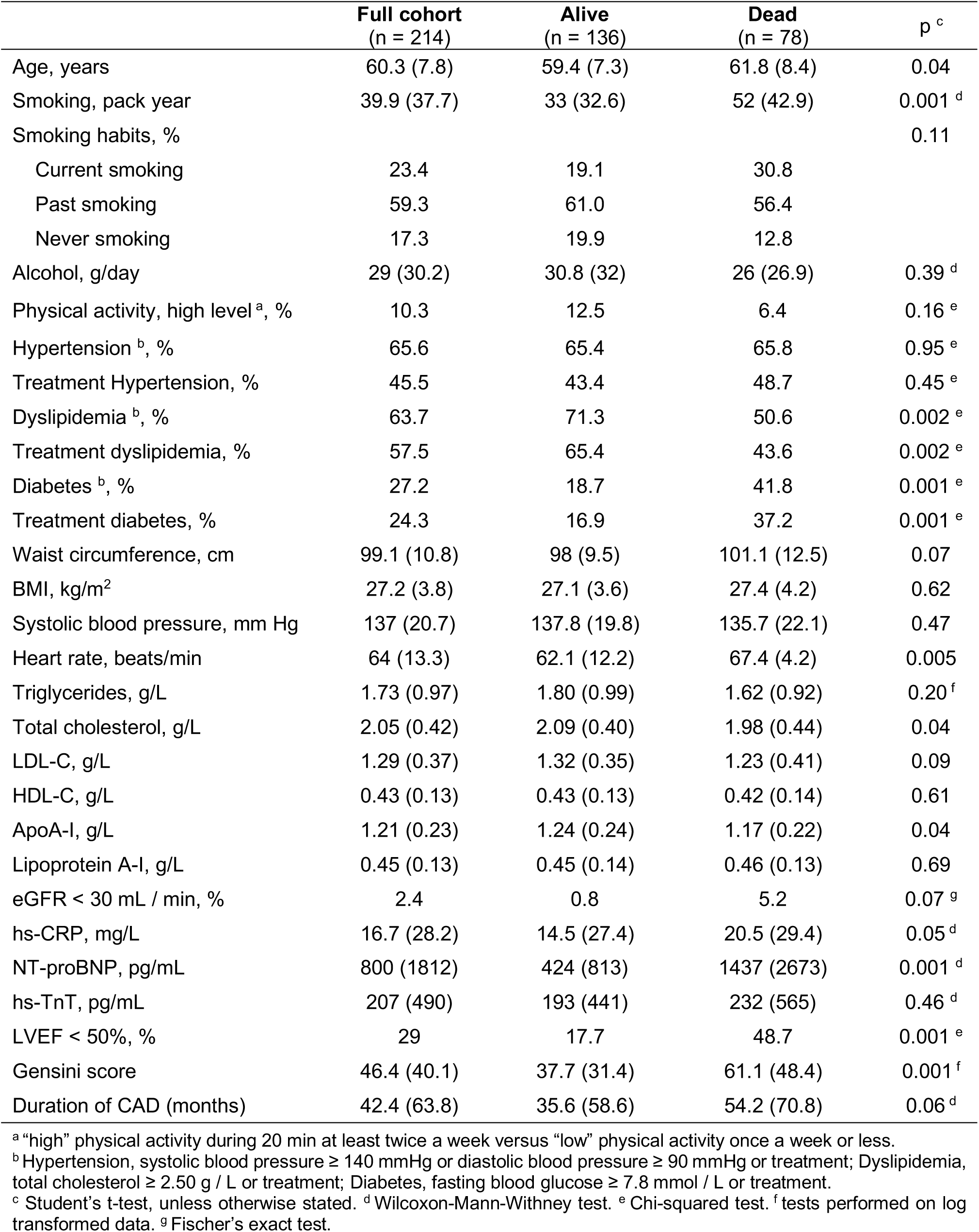
Clinical and biological characteristics in coronary artery disease patients when they were first included in the GENES cohort. Data are expressed in mean (SD) or % (n). BMI, body mass index; hs-CRP, high-sensitivity C-reactive protein; eGFR, estimated glomerular filtration rate; hs-TnT, high-sensitive cardiac troponin T; NT-ProBNP, N-terminal pro-brain natriuretic peptide; LVEF, left ventricular ejection fraction; CAD, coronary artery disease.

|  | Full cohort<br>(n = 214) | Alive<br>(n = 136) | Dead<br>(n = 78) | p<sup>c</sup> |
| --- | --- | --- | --- | --- |
| Age, years | 60.3 (7.8) | 59.4 (7.3) | 61.8 (8.4) | 0.04 |
| Smoking, pack year | 39.9 (37.7) | 33 (32.6) | 52 (42.9) | 0.001<sup>d</sup> |
| Smoking habits, % |  |  |  | 0.11 |
| Current smoking | 23.4 | 19.1 | 30.8 |  |
| Past smoking | 59.3 | 61.0 | 56.4 |  |
| Never smoking | 17.3 | 19.9 | 12.8 |  |
| Alcohol, g/day | 29 (30.2) | 30.8 (32) | 26 (26.9) | 0.39<sup>d</sup> |
| Physical activity, high level<sup>a</sup>, % | 10.3 | 12.5 | 6.4 | 0.16<sup>e</sup> |
| Hypertension<sup>b</sup>, % | 65.6 | 65.4 | 65.8 | 0.95<sup>e</sup> |
| Treatment Hypertension, % | 45.5 | 43.4 | 48.7 | 0.45<sup>e</sup> |
| Dyslipidemia<sup>b</sup>, % | 63.7 | 71.3 | 50.6 | 0.002<sup>e</sup> |
| Treatment dyslipidemia, % | 57.5 | 65.4 | 43.6 | 0.002<sup>e</sup> |
| Diabetes<sup>b</sup>, % | 27.2 | 18.7 | 41.8 | 0.001<sup>e</sup> |
| Treatment diabetes, % | 24.3 | 16.9 | 37.2 | 0.001<sup>e</sup> |
| Waist circumference, cm | 99.1 (10.8) | 98 (9.5) | 101.1 (12.5) | 0.07 |
| BMI, kg/m<sup>2</sup> | 27.2 (3.8) | 27.1 (3.6) | 27.4 (4.2) | 0.62 |
| Systolic blood pressure, mm Hg | 137 (20.7) | 137.8 (19.8) | 135.7 (22.1) | 0.47 |
| Heart rate, beats/min | 64 (13.3) | 62.1 (12.2) | 67.4 (4.2) | 0.005 |
| Triglycerides, g/L | 1.73 (0.97) | 1.80 (0.99) | 1.62 (0.92) | 0.20<sup>f</sup> |
| Total cholesterol, g/L | 2.05 (0.42) | 2.09 (0.40) | 1.98 (0.44) | 0.04 |
| LDL-C, g/L | 1.29 (0.37) | 1.32 (0.35) | 1.23 (0.41) | 0.09 |
| HDL-C, g/L | 0.43 (0.13) | 0.43 (0.13) | 0.42 (0.14) | 0.61 |
| ApoA-I, g/L | 1.21 (0.23) | 1.24 (0.24) | 1.17 (0.22) | 0.04 |
| Lipoprotein A-I, g/L | 0.45 (0.13) | 0.45 (0.14) | 0.46 (0.13) | 0.69 |
| eGFR < 30 mL / min, % | 2.4 | 0.8 | 5.2 | 0.07<sup>g</sup> |
| hs-CRP, mg/L | 16.7 (28.2) | 14.5 (27.4) | 20.5 (29.4) | 0.05<sup>d</sup> |
| NT-proBNP, pg/mL | 800 (1812) | 424 (813) | 1437 (2673) | 0.001<sup>d</sup> |
| hs-TnT, pg/mL | 207 (490) | 193 (441) | 232 (565) | 0.46<sup>d</sup> |
| LVEF < 50%, % | 29 | 17.7 | 48.7 | 0.001<sup>e</sup> |
| Gensini score | 46.4 (40.1) | 37.7 (31.4) | 61.1 (48.4) | 0.001<sup>f</sup> |
| Duration of CAD (months) | 42.4 (63.8) | 35.6 (58.6) | 54.2 (70.8) | 0.06<sup>d</sup> |

<sup>a</sup> “high” physical activity during 20 min at least twice a week versus “low” physical activity once a week or less.
<sup>b</sup> Hypertension, systolic blood pressure $\geq 140$ mmHg or diastolic blood pressure $\geq 90$ mmHg or treatment; Dyslipidemia, total cholesterol $\geq 2.50$ g / L or treatment; Diabetes, fasting blood glucose $\geq 7.8$ mmol / L or treatment.
<sup>c</sup> Student's t-test, unless otherwise stated. <sup>d</sup> Wilcoxon-Mann-Whitney test. <sup>e</sup> Chi-squared test. <sup>f</sup> tests performed on log transformed data. <sup>g</sup> Fischer's exact test.

Hypertension was diagnosed or treated in 65.6 % patients (45.5 % under treatment) with no difference between the deceased and alive groups (p = 0.95). Dyslipidemia was diagnosed or treated in 71.3 % (65.4 % treated) of the alive group, and in 50.6 % (43.6 % upon treatment) of the deceased group and the difference was statistically significant (p = 0.002). Among patients under lipid lowering therapy, 87 % were treated with statins. Diabetes was diagnosed in 18.7 % (16.9 % treated) of the alive group and in 41.8 % (37.2 % upon treatment) of the deceased group and the difference was statistically significant (p = 0.001).

Patients having deceased during the follow-up period had a longer duration of CAD, a decreased left ventricle ejection volume (LVEF), a higher heart rate and a more severe angiographic lesion score (Gensini). Regarding cardiovascular risk factors, smoking habits and treatment for diabetes were more frequent in the deceased group, whereas lipid-lowering therapy was less frequent. Among lipoprotein parameters, only apoA-I, a major HDL marker, was significantly lower in further deceased patients. Hs-CRP, an inflammatory marker, was lower in surviving than in deceasing patients. NT-proBNP levels were significantly higher in further deceased patients but no difference was observed in the concentration of hs-TnT between the two groups, indicating that deceased patients suffered from a more severe myocardial dysfunction rather than from a more extended myocardial necrosis.

The distribution of prior cardiovascular events in the study population is shown in Supplementary Table 1. The majority is represented by myocardial infarction (MI, 52.3 %), followed by revascularization procedures (37.9 %) and then by stable ischemic heart disease (IHD, 8%). Logically, among patients deceased during follow-up, past history of MI and IHD were more frequent than in alive patients.

### HDL particles according to vital status

HDL particles’ profile was determined by NMR spectroscopy, enabling to distinguish large HDL (LHDL-P, 8.8 − 13 nm) and small-sized HDL (SHDL-P, 7.3 − 8.7 nm) particles. The latter accounted for about ∼ 85 % of total HDL particles (HDL-P). HDL-P was ∼ 10% lower in deceased than in surviving patients (24.6 μmol/L [SD, 6.0] vs. 27.5 μmol/L [SD, 4.9], p = 0.001, Table 2). This difference was entirely due to a decreased number of SHDL-P, whereas number LHDL-P was not different according to the vital status (Table 2). The average size of total HDL particles (HDL size) was found higher in deceased patients (8.94 nm *versus* 8.82 nm, p = 0.014).

**Table 2.**
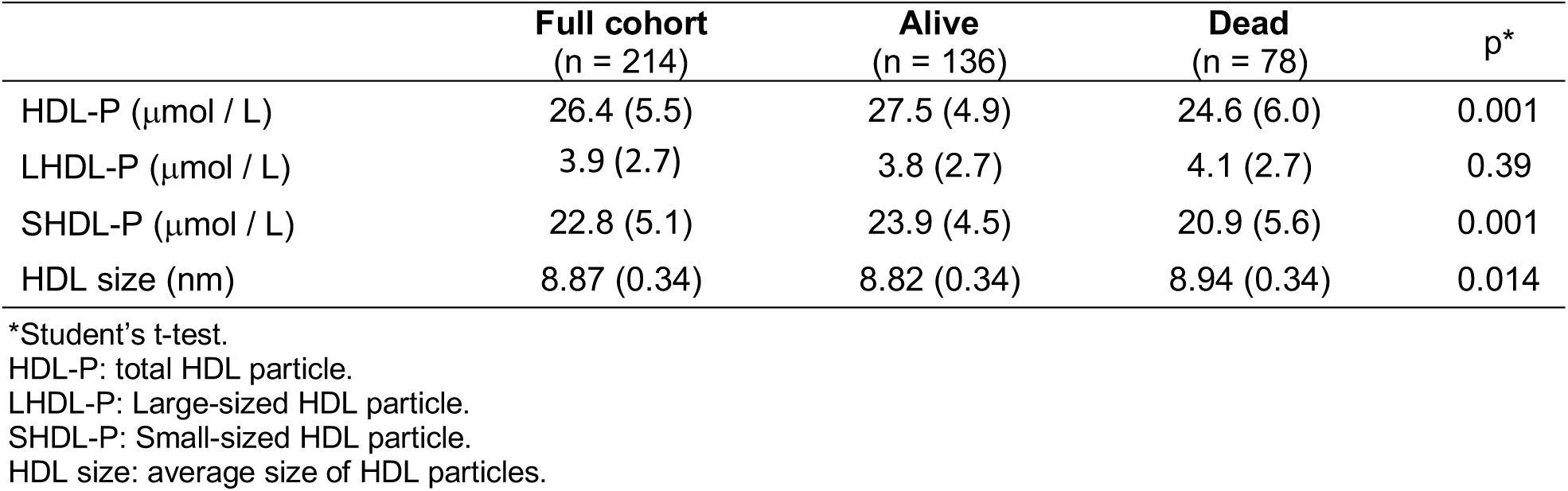
NMR HDL measures in coronary artery patients (n = 214) according to vital status. Data are expressed in mean (SD).

|  | **Full cohort**<br>(n = 214) | **Alive**<br>(n = 136) | **Dead**<br>(n = 78) | p* |
| --- | --- | --- | --- | --- |
| HDL-P (μmol / L) | 26.4 (5.5) | 27.5 (4.9) | 24.6 (6.0) | 0.001 |
| LHDL-P (μmol / L) | 3.9 (2.7) | 3.8 (2.7) | 4.1 (2.7) | 0.39 |
| SHDL-P (μmol / L) | 22.8 (5.1) | 23.9 (4.5) | 20.9 (5.6) | 0.001 |
| HDL size (nm) | 8.87 (0.34) | 8.82 (0.34) | 8.94 (0.34) | 0.014 |

\*Student's t-test.
HDL-P: total HDL particle.
LHDL-P: Large-sized HDL particle.
SHDL-P: Small-sized HDL particle.
HDL size: average size of HDL particles.

### Correlations between HDL particles measures and clinical and biological parameters

Correlations were investigated between markers of HDL particles and other clinical or biological parameters in the study population (Table 3). All NMR-HDL measures were correlated between themselves and with other HDL markers: HDL-C, apoA-I and lipoprotein A-I (LpA-I). Logically, the average HDL size was correlated positively with the number of LHDL-P, and negatively, with the number of SHDL-P. Triglycerides were associated positively with HDL-P and SHDL-P but negatively with LHDL-P, mean HDL size and HDL-C. No association was observed with apo A-I. These correlations might reflect the remodeling of HDL lipids induced by the cholesterol ester transfer protein (CETP) acting between HDL particles and triglyceride-rich lipoproteins. Alcohol consumption positively correlated with HDL-C and HDL-P, and more specifically with SHDL-P but not with apoA-I. Inflammation, as documented by plasma hs-CRP, was inversely associated with apo A-I, HDL-P and SHDL-P, but not with LHDL-P. HDL-P, SHDL-P and apo A-I were negatively associated with NT-proBNP and hs-TnT

**Table 3.**
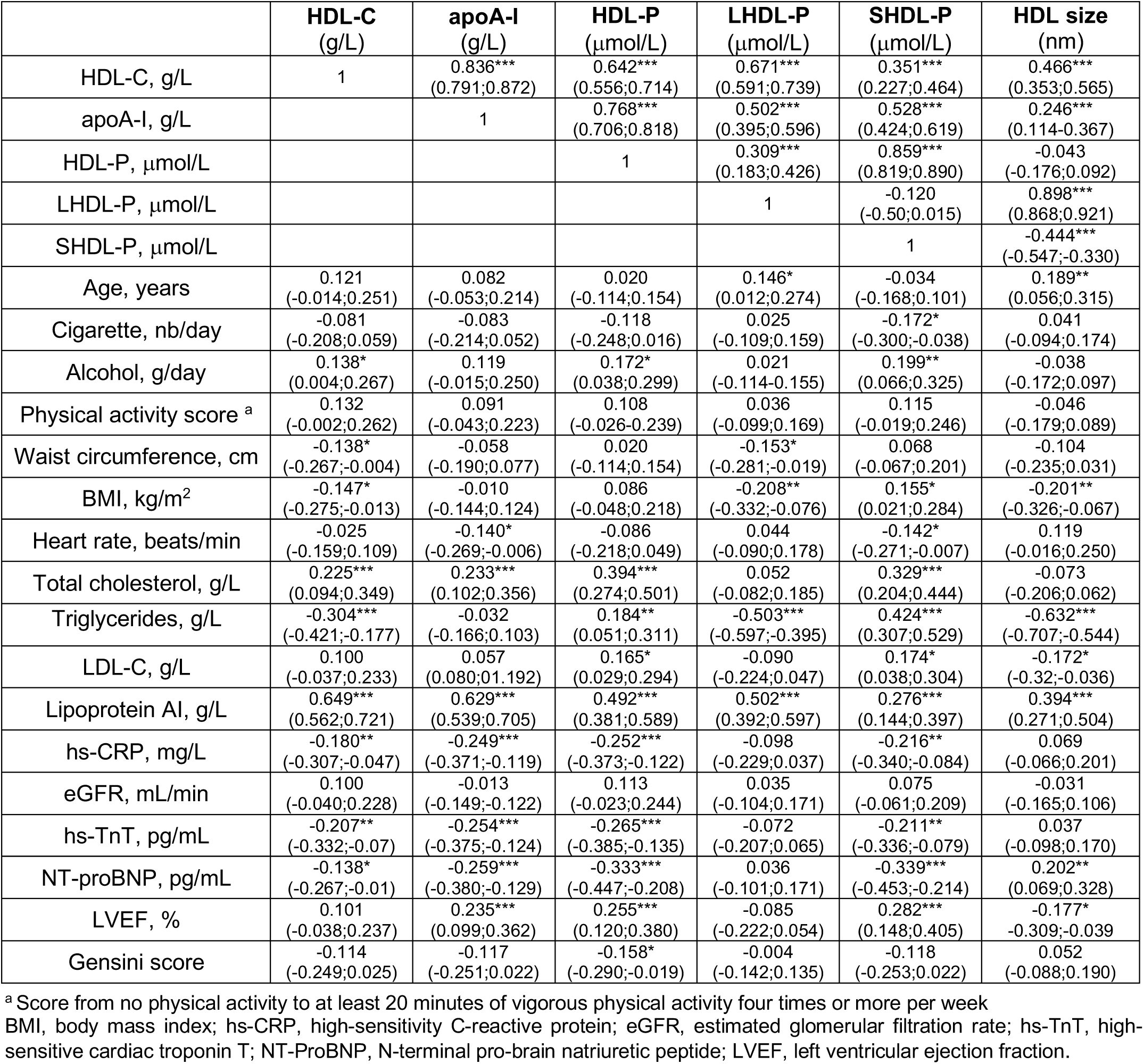
Correlation between NMR HDL measure and biological and biochemical parameters and other cardiovascular risk factors in coronary artery disease patients. Spearman rank correlation coefficients (95% confidence interval). *p<0.05, ** p<0.01, *** p<0.001

The severity of coronary lesions, as illustrated by the Gensini score, was inversely related to HDL-P. Strong positive associations were observed between LVEF and both HDL-P and SHDL-P. ApoA-I levels correlated with LVEF but not significantly with the Gensini score. For comparison, no relationship was recorded between HDL-C and either LVEF or the Gensini score. Thus, although HDL markers were strongly correlated between themselves, total and subclasses of HDL particles displayed specific association with clinical variables.

### Total and cardiovascular mortality according to tertiles of HDL markers

Each one HDL marker was considered according to tertiles of its distribution in the whole study population (Table 4). Death rates during follow-up were determined across the different tertiles and associations were determined after adjustment on classical risk factors. Similar associations were observed without adjustment. The strongest association to total and cardiovascular mortality was observed for HDL-P distribution. A 45% reduction in death rates was recorded in tertiles 2 and 3, as compared to tertile 1. Each 1 SD increase in HDL particles number was found associated with ∼ 42 % reduction in total or cardiovascular mortality (HR = 0.58 [95%CI, 0.45-0.75] and 0.59 [95%CI, 0.44-0.80], respectively). Results were almost identical considering SHDL-P. By contrast, no association between LHDL-P and mortality was observed, except for an almost significant positive trend (p = 0.07) between LHDL-P and death rates. Concordantly, death rates were significantly different across HDL size distribution, an increase in particles size being associated with highest death rates. Considering the “classical” HDL markers, HDL-C tertiles did not display different death rates. On the other hand, apoA-I distribution was associated to total and cardiovascular mortality; each 1 standard deviation increase of apoA-I was associated to a ∼ 31% risk reduction (HR = 0.69 [95%CI, 0.54-0.88] and 0.69 [95%CI, 0.49-0.91], respectively).

**Table 4.**
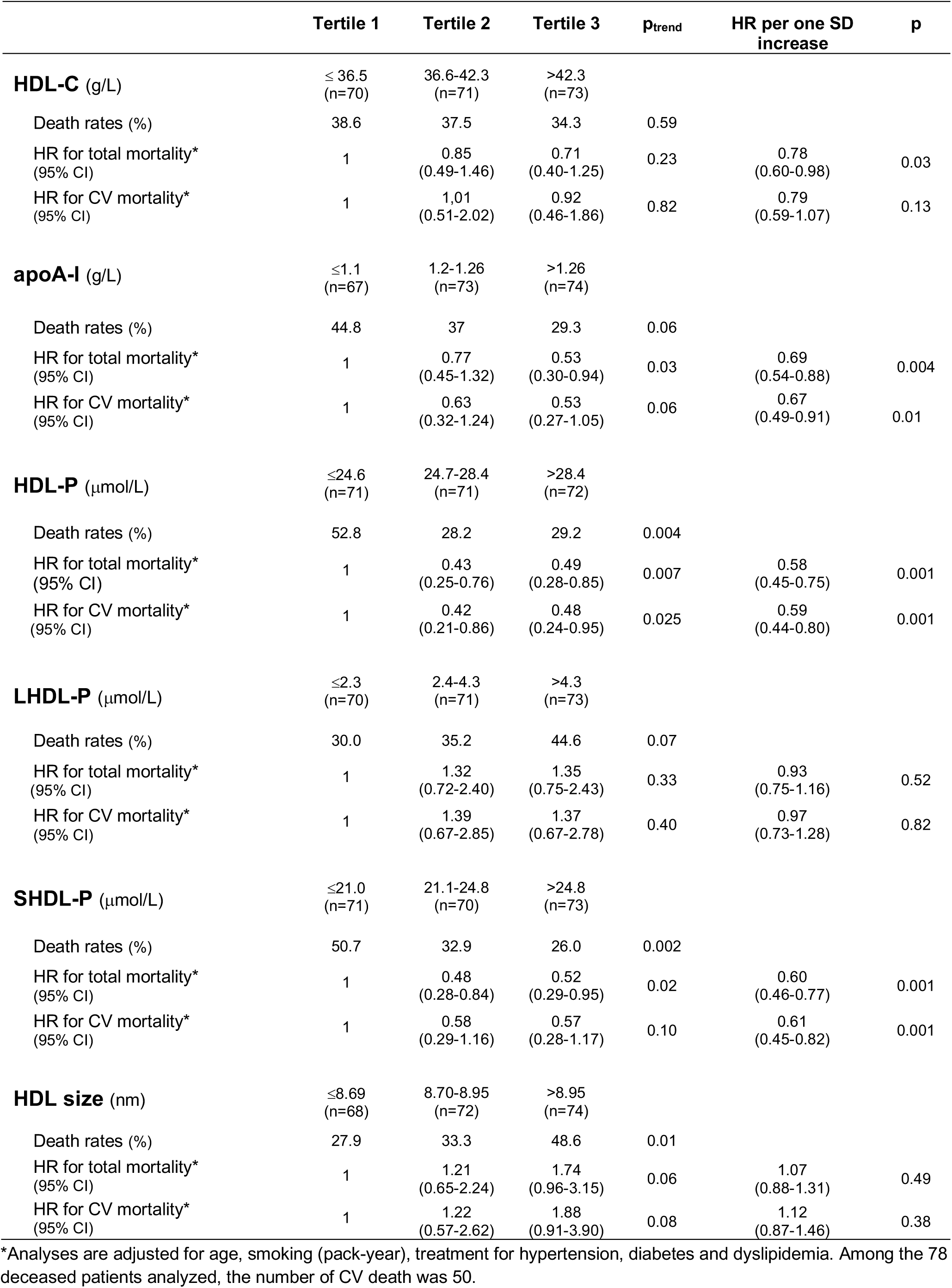
Death rate according to tertiles of HDL-related biomarkers and association with total and cardiovascular (CV) mortality. HR: Hazard Ratio; CI: Confidence Interval.

A further multivariate analysis was conducted, including adjustment on an extended panel of bio-clinical variables reflecting cardiovascular risk factors, including BMI and physical activity, smoking, alcohol consumption, treatments (diabetes, hypertension, dyslipidemia), heart condition including heart rate, CAD duration, Gensini score and LVEF, renal function (eGFR) and inflammation (hs-CRP). Hazard ratios for all-cause mortality per 1-SD increase of HDL-P, of SHDL-P or of apoA-I were all significant. They were lower for HDL-P (HR = 0.58 [95%CI, 0.43-0.77]) and SHDL-P (HR = 0.57 [95%CI, 0.42-0.77]) than for apoA-I (HR = 0.68 [95%CI, 0.51-0.90]) (Figure 1). Almost identical data were obtained for cardiovascular mortality (Figure 1, model 1). Further adjustment for non-HDL-C did not diminish the inverse association of apoA-I, HDL-P and SHDL-P with all-cause of mortality. Those associations were also maintained for cardiovascular mortality, except with SHDL-P (p = 0.060). We have also adjusted for cardiac markers, NT-ProBNP and hs-TnT (Figure 1, model 3), and all associations were maintained.

**Figure 1.**
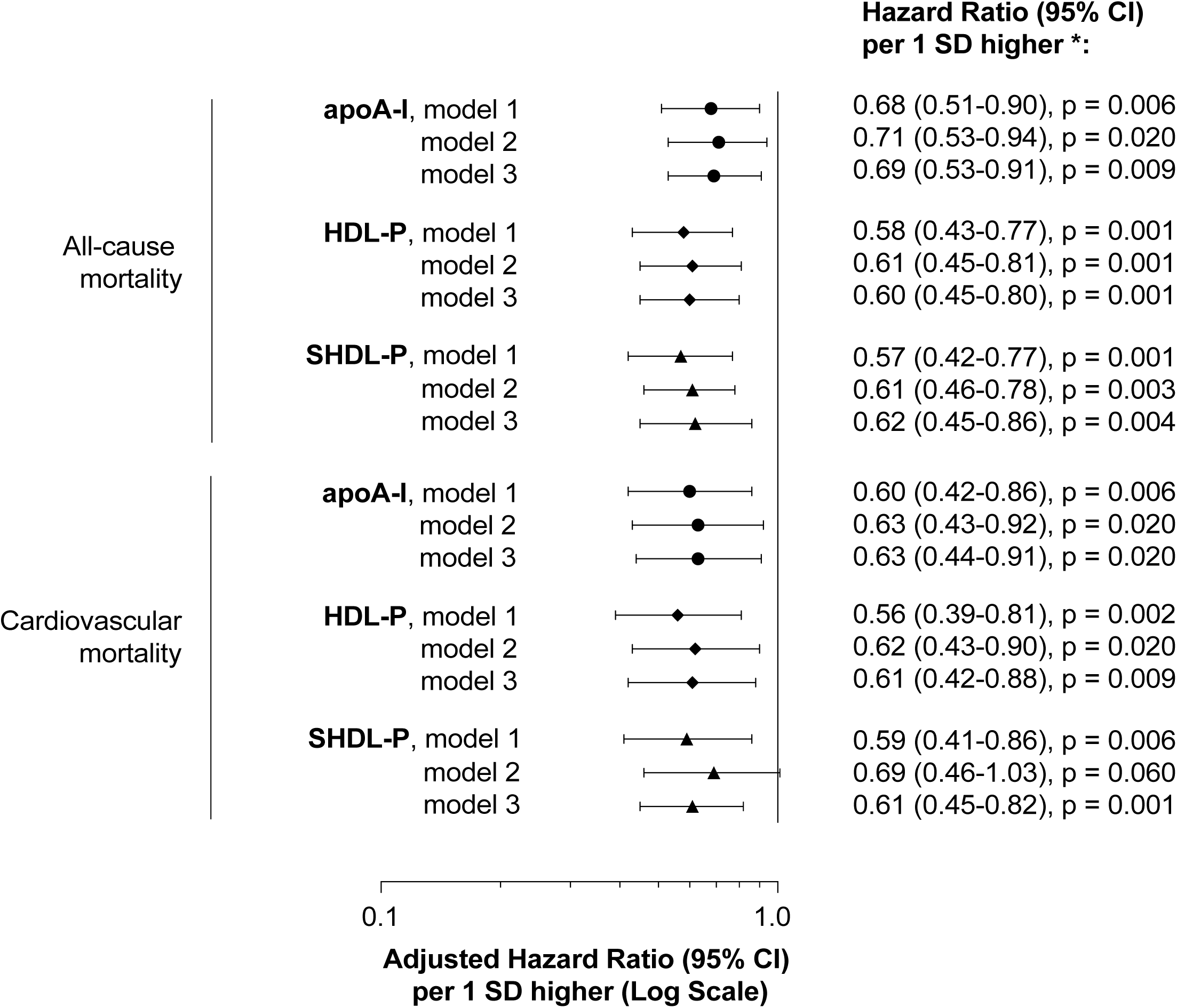
Relative risk of all-cause and cardiovascular mortality as a function of apoA-I, HDL-P and SHDL-P. Graphic represents hazard ratios (dots) and corresponding 95% confidence interval (95%CI) for risk of all-cause and cardiovascular mortality per 1 standard deviation increase of apoA-I, HDL-P or SHDL-P. Among the 78 deceased patients analyzed, the number of CV death was 50. Model 1: adjusted for age, smoking (pack-year), alcohol consumption, physical activity, BMI, treatments for dyslipidemia, hypertension and diabetes, hs-CRP, eGFR, heart rate, LVEF, duration of CAD and Gensini score. Model 2: Model 1 plus non-HDL-C. Model 3: Model 1 plus NT-proBNP and hs-TnT.

Associations between HDL markers and mortality are illustrated in the survival curves established during the whole follow-up period for the different tertiles (Figure 2). Death rates were regular during the whole time course of follow-up. For both HDL-P and SHDL-P, patients in the first tertile had a poorer survival than patients in tertiles 2 and 3. This was particularly evident for early events: of the 17 deaths recorded during the 0-3 year follow-up, 11 (65%) were in the lowest tertile of HDL-P as compared to 8 out of 20 (40%) for the deaths recorded in the 9-12 year period. A comparable trend was observed for apoA-I distribution yet survival differences during the whole period did not reach statistical significance.

**Figure 2.**
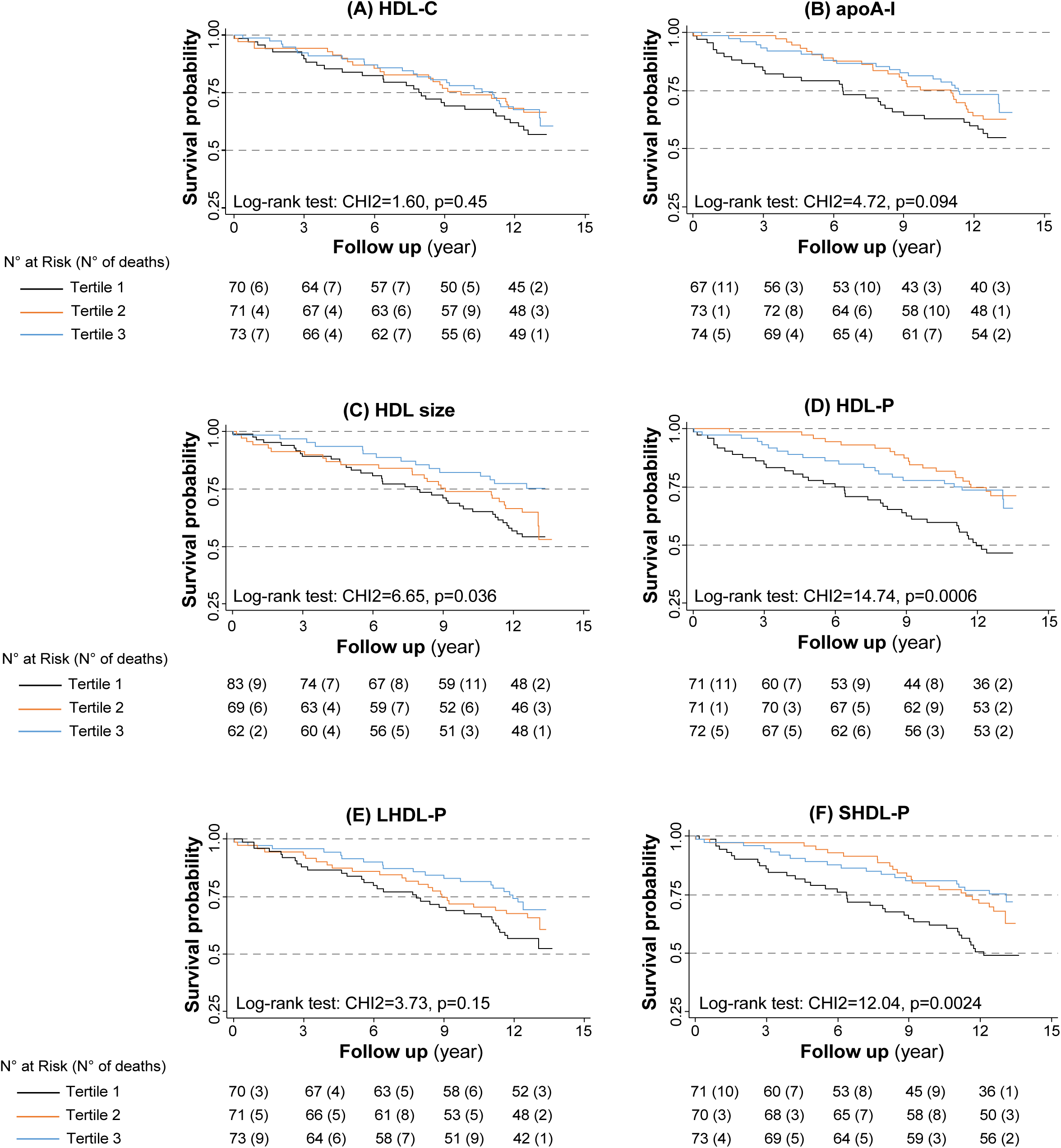
Kaplan-Meier survival curves. Survival curves for the follow up period were established as a function of HDL-C (A), apoA-I (B), HDL size (C), HDL-P (D), LHDL-P (E) and SHDL-P (F) tertiles. Tertiles range for HDL-C: ≤ 36.5, 36.6 – 42.3, > 42.3 g/L; tertiles range for apoA-I: ≤ 1.1, 1.2 – 1.26, > 1.26 g/L; tertiles range for HDL size: ≤ 8.69, 8.70 – 8.95, > 8.95 g/L; tertiles range for HDL-P: ≤ 24.6, 24.7 – 28.4, > 28.4 μmol/L; tertiles range for LHDL-P: ≤ 2.3, 2.4 – 4.3, > 4.3 μmol/L; tertile for SHDL-P: ≤ 21.0, 21.1 – 24.8, > 24.8 μmol/L

## Discussion

In the present study, serum levels of total HDL particles (HDL-P) and of small-sized HDL particles (SHDL-P) were inversely related to all-cause as well as to specific cardiovascular mortality in CAD patients. Every 1-SD increase of HDL particle number was associated to a 44% decrease in cardiovascular mortality, after multiple adjustments on cardiovascular risk factors and on clinical markers of heart condition. Among other HDL markers, apoA-I was also inversely related, though to a lesser extent, to total and cardiovascular mortality. Conversely, HDL-C or large HDL particles (LHDL-P) were not associated with mortality. However, higher death rates were recorded as average HDL particle size increased.

NMR-based lipoprotein profiling methods are not yet standardized and the estimated diameter range of HDL subclasses depends on the analytical methods used. Also, no classification for HDL subclasses analysed by NMR has been yet approved for routine purpose. For instance, the analytical method developed by LipoScience (now LabCorp, Raleigh, NC) grouped HDL particles into 3 subclasses from small (S-HDL-P, 7.3 − 8.2 nm), medium (M-HDL-P, 8.2 − 9.4 nm) to large (L-HDL-P, 9.4 − 14 nm) with small and medium-sized HDL particles being sometimes grouped (MS-HDL-P, 7.3 − 9.4 nm), while the AXINON^®^ lipoFIT^®^-S100 system (Numares AG, Regensburg, Germany) used in the present study grouped HDL subclasses into small (SHDL-P, 7.3 − 8.7 nm) and large (LHDL-P, 8.8 − 13 nm)^16,18^. In this latest classification, it is worth mentioning that the estimated diameter ranges for SHDL-P and LHDL-P were similar to those for HDL_3c + 3b + 3a_ (7.2 − 7.8 nm; 7.8 − 8.2 nm; 8.2− 8.8 nm) and HDL_2a + 2b_ (8.8 −9.7 nm; 9.7 − 12.9 nm), respectively, as isolated from plasma by density gradient ultracentrifugation and analyzed by electrophoresis on a nondenaturing gel^19,20^.

To date, the inverse and independent association between HDL-P / MS-HDL-P and cardiovascular risk has been extensively documented with respect to primary prevention, either in individuals without baseline CAD^5,21^ and in those with pre-clinical atherosclerosis, as documented by carotid intima-media thickness^5,22^, or coronary calcifications^23^. More recently, in a large study carried out in high-risk individuals undergoing coronary catheterization for suspicion of CAD, followed-up during 8 years, HDL-P and MS-HDL-P were independent predictors of all-cause mortality^6^. With respect to secondary prevention, only a few studies have evaluated the relationship of NMR-derived HDL particle subclasses with cardiovascular disease outcomes in CAD patients. In a prospective nested case-control study of 364 men with new CAD events (non-fatal myocardial infarction or cardiac death) during a 5.1 year follow-up paired to 697 age-matched control, total HDL-P and small HDL-P (7.3 − 8.2 nm) were strong, independent, predictors of recurrent coronary events, whereas levels of HDL-C were not^24^. In patients suffering from acute heart failure, concentrations of both HDL-P and small HDL-P (7.3 − 8.7 nm) were inversely related to short-term (3-month) mortality, after multiple adjustments on confounding variables, including NT-proBNP, a classical marker of heart failure^17^.

The present study brings some additional insights on this relation between NMR-derived HDL particles subclasses and long-term prognosis in CAD patients, with angiographically documented coronary lesions. Association of HDL markers to all-cause and cardiovascular mortality was assessed after adjustment on a large variety of confounders: life-style parameters, clinical and biological variables documenting cardiovascular risk factors, inflammatory status, renal function and heart condition. In addition, this study has enabled to compare the predictive value of apoA-I versus HDL-P measurements, which showed comparable associations, although Hazard ratios were better with HDL-P levels.

HDL particles, and most particularly small-sized HDL, may act against atherosclerosis through different mechanisms. Small HDL behave as the best acceptors of ABCA1-mediated cholesterol efflux from macrophages, leading subsequently to the mobilization of intracellular cholesterol to the plasma membrane^25,26^. Small and dense HDL particles also protect LDL from oxidation. HDL particles act through removing phospholipid hydroperoxides from LDL and by inactivating oxidized lipids by specific enzymes like paraoxonase-1 (PON-1) and PAF-acetylhydrolase^27,28^. Moreover small protein-rich HDL exert anti-inflammatory properties by depressing expression of VCAM-1 at the surface of endothelial cells^29^. On these cells, HDL particles appear to be cytoprotective by inhibiting apoptosis induced by oxidized LDL, and small HDL_3_ would be the most effective in this function^30^. Altogether those observations suggest that the proteome associated to small HDL particles support various biological activities, which impair atherosclerosis development.

Moreover, HDL particles may exert beneficial effects on myocardial functions. Indeed, in different experimental contexts, it was demonstrated that HDL particles protect against ischemia reperfusion injury^31^, leading to a reduction in infarct size. HDL may also improve myocardial function by reducing ventricular remodelling following infarction^32^. In isolated cardiomyocytes, HDL particles were shown to prevent apoptosis through an AMP-kinase dependent mechanism^33^. These experimental observations on a direct impact of HDL on myocardial functions might translate into clinical impacts. In support of this concept is the positive correlation observed here between HDL-P, small HDL-P and the left ventricular ejection fraction, concordant with the negative association between HDL-P, small HDL-P and NT-proBNP observed here and previously reported^17^.

In this study, concentrations of large HDL particles were not associated to mortality. However, higher death rates were recorded as HDL size increased (p < 0.01); following multiple adjustments, association to total mortality for the upper tertile of HDL size was close to statistical significance (p = 0.06). Similar observations regarding all-cause mortality in individuals who are at high cardiovascular risk have been previously reported^6^. Large HDL-P might be less effective than SHDL-P regarding various atheroprotective functions, like cholesterol efflux, anti-oxidative and anti-inflammatory properties, and cytoprotective effects on endothelium^4,34^. Moreover, accumulation of large HDL might reflect a defect in HDL catabolism, and particularly in HDL liver uptake, which constitutes the last step of reverse cholesterol transport^35^. Similarly, we did not observe any association of HDL-C with mortality. This is concordant with the lack of association between LHDL-P and mortality, since HDL-C mainly reflects cholesterol associated with large, lipid rich, HDL particles.

ApoA-I was inversely related to mortality: for each 1-SD increase of apoA-I, a 31% and 33% decrease in all-cause and cardiovascular mortality was recorded, respectively. So far, apoA-I has been little used in epidemiological studies. However, calibration on reference international standards has made the immunoassay of apoA-I robust and comparable between studies. Furthermore, apoA-I measurement is much less influenced than HDL-C by intravascular enzymes and lipid transfer proteins, which participate in HDL remodelling. Thus, apoA-I measurement may improve assessment of cardiovascular risk^36^. Association to mortality was somewhat weaker for apoA-I than for HDL-P or SHDL-P. This might be explained by the fact that the apoA-I content per particle varies on average from 2 to 4, between small HDL_3_ and large HDL_2_^37^, so that large HDL particles are somewhat overrepresented in apoA-I quantification.

A number of limitations of the present study must be noted. First, the small size of the study is a limitation. Indeed, the first 214 consecutive CAD patients of the CAD patients from the GENES cohort where included in the present study, which represent 25 % of the whole cohort (n = 834). Accordingly, mortality rate was higher in included patients than in the whole cohort (36.4% *versus* 29.1%, data not shown). However, similar clinical and biological characteristics were measured between included and non-included patients (Supplementary Table 2). Second, this study was designed only with men, which has the advantage of recording a larger number of events than in a mixed all-gender cohort, but limited the translatability of our results to women. Finally, despite adjustments on established CVD risk factors the possibility remains that other important confounding variables with effects on HDL parameters were not measured or considered in our analyses.

In conclusion, the present study demonstrated that the concentration of total HDL particles and small-sized HDL particles may serve as a better prediction tool than HDL-C and apoA-I to assess long term prognosis in coronary patients. Studies on HDL metabolism had progressively led to the schematic view of an interconversion cycle of HDL particles in the plasma compartment, driven by cell cholesterol efflux, enzymes like LCAT, lipases and lipid transfer proteins^38,39^. More recently the concept has emerged that HDL particles of different geometry and chemical composition have distinct metabolic fate and display specific functional properties^3,4,40^. This supports the idea that HDL functionality might be more precisely assessed by the quantification of specific HDL particles with high atheroprotective effects. In the future, quantification of HDL particle concentration and HDL subclasses along with HDL functional measurement could be clinically useful in CVD risk assessment.

## Acknowledgments

This work was supported by the French National Research Agency (ANR, #ANR-16-CE18-0014-01), European Regional Development Fund (ERDF) and “La Région Occitanie” (Project THERANOVASC n° ESR_R&S_DF-000094 / 2018-003303 / 18009464)

## Abbreviations

apoA-I: Apolipoprotein A-I
BMI: Body Mass Index
CAD: Coronary artery disease
CI: Confidence interval
eGFR: Estimated Glomerular Filtration Rate
HDL: High-density lipoprotein
HDL-P: HDL particle
LHDL-P: Large-sized HDL-particle
SHDL-P: Small-sized HDL-particle
HDL size: Average HDL particle size.
HR: Hazard Ratio
hs-CRP: High-sensitivity C-Reactive Protein
LDL: Low-density lipoprotein
LpA-I: Lipoprotein A-I
LVEF: Left Ventricular Ejection Fraction
SD: Standard Deviation

