## Supplemental Table 1 for "Serum level of High-density lipoprotein particles are independently associated with long-term prognosis in patients with coronary artery disease: The GENES study"

| | Full cohort (n = 214) | Alive (n = 136) | Dead (n = 78) | p |
|---|---|---|---|---|
| | | | | <0.02 |
| Myocardial infarction (MI) | 112 (52.3%) | 66 (48.5 %) | 46 (59.0 %) | |
| Ischemic heart disease (IHD) | 21 (9.8%) | 9 (6.6 %) | 12 (15.4%) | |
| Revascularization | 81 (37.9%) | 61 (44.9 %) | 20 (25.6%) | |

**Supplementary Table 1. Distribution of past history events**

Sorted in the following order: first MI even if they had revascularization then IHD then revascularization
