## Supplemental Table 2 for "Serum level of High-density lipoprotein particles are independently associated with long-term prognosis in patients with coronary artery disease: The GENES study"

| | Included (n = 214) | Non included (n = 620) | p [c] |
|---|---|---|---|
| Age, years | 60.3 (7.8) | 60.1 (8.2) | 0.81 |
| Smoking, pack year | 39.9 (37.7) | 36.9 (35.3) | 0.29 [d] |
| Smoking habits, % | | | 0.09 |
| Current smoking | 23.4 | 18.9 | |
| Past smoking | 59.3 | 63.5 | |
| Never smoking | 17.3 | 17.6 | |
| Alcohol, g/day | 29 (30.2) | 28 (32.6) | 0.75 [d] |
| Physical activity, high level [a], % | 10.3 | 12.6 | 0.46 [e] |
| Hypertension [b], % | 65.6 | 69.2 | 0.33 [e] |
| Treatment Hypertension, % | 45.5 | 43.4 | 0.66 [e] |
| Dyslipidemia [b], % | 63.7 | 71.1 | 0.05 [e] |
| Treatment dyslipidemia, % | 57.5 | 65.8 | 0.04 [e] |
| Diabetes [b], % | 27.2 | 27.2 | 0.99 [e] |
| Treatment diabetes, % | 24.3 | 24.5 | 0.92 [e] |
| Waist circumference, cm | 99.1 (10.8) | 98.9 (11.0) | 0.82 |
| BMI, kg/m$^2$ | 27.2 (3.8) | 27.4 (4.1) | 0.34 |
| Systolic blood pressure, mm Hg | 137 (20.7) | 140.5 (20.4) | 0.03 |
| Heart rate, beats/min | 64 (13.3) | 64.2 (11.6) | 0.81 |
| Triglycerides, g/L | 1.73 (0.97) | 1.70 (1.02) | 0.66 [f] |
| Total cholesterol, g/L | 2.05 (0.42) | 2.00 (0.44) | 0.56 |
| LDL-C, g/L | 1.29 (0.37) | 1.23 (0.39) | 0.09 |
| HDL-C, g/L | 0.43 (0.13) | 0.43 (0.12) | 0.41 |
| ApoA-I, g/L | 1.21 (0.23) | 1.25 (0.22) | 0.06 |
| Lipoprotein A-I, g/L | 0.45 (0.13) | 0.48 (0.15) | 0.06 |
| eGFR < 30 mL / min, % | 2.4 | 1.5 | 0.37 [g] |
| hs-CRP, mg/L | 16.7 (28.2) | 11.6 (19.0) | 0.001 [d] |
| NT-proBNP, pg/mL | 800 (1812) | 607 (1433) | 0.23 [d] |
| hs-TnT, pg/mL | 207 (490) | 123 (342) | 0.02 [d] |
| LVEF < 50%, % | 29 | 26.1 | 0.37 [e] |
| Gensini score | 46.4 (40.1) | 47.2 (40.5) | 0.89 [f] |
| Duration of CAD (months) | 42.4 (63.8) | 42.7 (66.6) | 0.92 [d] |

**Supplementary Table 2. Comparison between CAD patients included and patients non included.**

Data are expressed in mean (SD) or % (n).

BMI, body mass index; hs-CRP, high-sensitivity C-reactive protein; eGFR, estimated glomerular filtration rate; hs-TnT, high-sensitive cardiac troponin T; NT-ProBNP, N-terminal pro-brain natriuretic peptide; LVEF, left ventricular ejection fraction; CAD, coronary artery disease.

[a] "high" physical activity during 20 min at least twice a week versus "low" physical activity once a week or less. [b] Hypertension, systolic blood pressure ≥ 140 mmHg or diastolic blood pressure ≥ 90 mmHg or treatment; Dyslipidemia, total cholesterol ≥ 2.50 g / L or treatment; Diabetes, glucose ≥ 7.8 mmol / L or treatment. [c] Student's t-test, unless otherwise stated. [d] Wilcoxon-Mann-Withney test; [e] Chi-squared test. [f] tests performed on log transformed data. [g] Fischer's exact test.
